# Surface residues and non-additive interactions stabilize a consensus homeodomain protein

**DOI:** 10.1101/2021.04.19.440332

**Authors:** Matt Sternke, Katherine W. Tripp, Doug Barrick

## Abstract

Despite the widely reported success of consensus design in producing highly stabilized proteins, little is known about the physical mechanisms underlying this stabilization. Here we explore the potential sources of stabilization by performing a systematic analysis of the 29 substitutions that we previously found to collectively stabilize a consensus homeodomain compared to an extant homeodomain. By separately introducing groups of consensus substitutions that alter or preserve charge state, occur at varying degrees of residue burial, and occur at positions of varying degrees of conservation, we determine the extent to which these three features contribute to the consensus stability enhancement. Surprisingly, we find that the largest total contribution to stability comes from consensus substitutions on the protein surface and that the largest per-substitution contributions come from substitutions that maintain charge state, suggesting that although consensus proteins are often enriched in charged residues, consensus stabilization does not result primarily from charge-charge interactions. Although consensus substitutions at strongly conserved positions also contribute disproportionately to stabilization, significant stabilization is also contributed from substitutions at weakly conserved positions. Furthermore, we find that identical consensus substitutions show larger stabilizing effects when introduced into the consensus background than when introduced into an extant homeodomain, indicating that synergistic, stabilizing interactions among the consensus residues contribute to consensus stability enhancement of the homeodomain.

**Significance Statement:** Proteins composed of consensus sequences from multiple sequence alignments are often more stable than extant proteins used to create them. Often about half the residues in a consensus protein differ from those of extant proteins. The contributions of these differences to stability are unknown. Here we substitute groups of residues with different properties (conservation, charge variation, solvent accessibility) to determine which substitutions lead to consensus stabilization. We find that surface and charge-conserving substitutions contribute to stability, that weakly-conserved substitutions make a significant collective contribution to stability, and that there is a significant non-additive contribution to stability in the consensus background. These results provide insights to the sequence origins of consensus stabilization and the evolutionary constraints that determine protein sequences.

## Introduction

Most natural proteins have stabilities ranging from 5-15 kcal mol^-1^ (1). These modest stabilities have been argued to result from weak selective pressure on stability, rather than an upper limit to stability (1, 2). Indeed, the stabilities of proteins from thermophilic organisms (3) and non-natural designed proteins (4) significantly exceed this marginal range. Furthermore, natural protein sequences can be stabilized by point substitutions (5, 6). Thus, there is great potential to design novel highly stable proteins as well as stabilize natural protein folds beyond their marginal stabilities.

Designing protein for high stability, however, has proven to be a significant challenge, since most substitutions are destabilizing (5, 6). Although a number stabilized proteins have been created using computational and rational design (4, 7–9) as well as laboratory evolution methods (10, 11), success rates can be low and methods are often labor-intensive and require considerable expertise. As one indication of the challenges that persist in protein engineering, state-of-the-art computational methods often show limited predictive power in the simple task of correctly classifying point substitutions as stabilizing versus destabilizing (12).

The rapid growth of protein sequence databases over the last decade has enabled sequence-based methods in protein design (13, 14). One sequence-based method, consensus sequence design, has shown considerable promise in generating highly stabilized natural protein folds. In consensus design, a consensus protein is generated in which the residue at each position is the most frequent residue in a corresponding multiple sequence alignment (MSA). Consensus design has been shown to stabilize proteins across a number of protein families with different architectures, chain lengths, phylogenetic distributions, and functions in most (but not all) cases (13, 15).

Despite the broadly demonstrated success of consensus design in increasing protein stability, little is known of the sequence, structural, or physical origins of consensus stabilization. Rather than optimizing protein sequences based on physical principles, consensus design is a bioinformatic approach. The underlying physical basis for consensus stabilization is encoded with the conservation of residues in an MSA. One general sequence feature that distinguish consensus proteins from natural proteins is an increase in the number of charged residues at the expense of polar uncharged residues (16). Differences between consensus sequences and natural sequences occur most frequently at positions of low conservation and positions on the protein surface (16). Identifying which of these general classes of substitutions gives rise to consensus stabilization may allow for further stabilization, and provide a route to achieve maximal stability with a minimal number of substitutions.

Surprisingly, although full consensus proteins are typically more stable than extant sequences from which they derive, individual consensus substitutions in extant proteins are often destabilizing. Only about half of substitutions toward consensus are stabilizing; the other half are destabilizing or have no effect on stability (13). An additive model in which the effect on stability of the full consensus protein is the sum of effects from the individual substitutions predicts a minimal increase in stability from full consensus substitution since the stabilizing substitutions would be offset by the destabilizing substitutions (17). This discrepancy suggests non-additive interactions among consensus residues may contribute to b consensus stabilization, however such interactions have not been reported in consensus proteins.

To determine which types of substitutions give rise to consensus stability enhancement, and to probe non-additive stabilization among consensus residues, we performed a mutational analysis using a consensus homeodomain (CHD) that we previously found to be stabilized by ∼4.5 kcal mol^-1^ relative to the well-studied *Drosophila melanogaster* Engrailed homeodomain (EnHD) (18). To directly determine the contributions to stability from specific sequence and structural features, we introduced sets of consensus substitutions that share a specific feature into EnHD, and characterized the thermodynamic stabilities of the resulting variants. These features include residue charge state, degree of side-chain burial, and extent of conservation. In addition, we probed non-additivity by comparing the effect of identical consensus substitutions (both larger sets of residues and individual residues) in the EnHD and CHD backgrounds. We find that the largest total contributions to stability arise from consensus substitutions on the protein surface, while the largest per-residue stabilizing effects come from substitutions that maintain charge state and substitutions at strongly conserved positions. However, no single feature accounts for the entire consensus stability increment. We also find that stabilizing effects correlate with the number of consensus substitutions introduced in each variant, suggesting that consensus stabilization arises from marginal, incremental effects distributed across each feature.

Furthermore, we find that some consensus residue substitutions show larger stabilizing effects when introduced in the CHD background than the EnHD background, suggesting that favorable non-additive couplings among consensus residues contribute to the significantly enhanced stability of the consensus homeodomain.

## Materials and methods

### Cloning, protein expression and purification

Genes encoding EnHD and CHD in pET24 expression plasmids were described previously.(18) Single residue substitutions were introduced using a Quikchange site-directed mutagenesis kit (Agilent). Genes encoding sequences with multiple substitutions were synthesized by GeneArt (ThermoFisher Scientific) and cloned into the pET24 expression plasmids using a Gibson Assembly Mastermix (New England Biolabs). All constructs contain an N-terminal Met-Gly-Ser and a C-terminal His_6_ tag. Proteins were expressed and purified as previously described (18).

### Equilibrium guanidine hydrochloride (GdnHCl) denaturations

Equilibrium GdnHCl-induced folding/unfolding transitions were monitored by circular dichroism (CD) using an Aviv model 400 spectropolarimeter. All experiments were performed using a Hamilton automated titrator. For each experiment, samples with identical protein concentrations were prepared in a native buffer containing 25 mM NaPO_4_ (pH 7.0) and 150 mM NaCl and a denaturing buffer containing 25 mM NaPO_4_ (pH 7.0) and 150 mM NaCl with a high concentration of GdnHCl (∼8 M for most experiments). GdnHCl concentrations in denatured protein solutions were determined using refractometry (19). Because the proteins were found to fold and unfold reversibly, we collected denaturation curves in either the forward (titrating in the denaturant) or reverse direction (diluting away the denaturant) to optimize sampling of the folded and unfolded baselines. For proteins of lower stability (for example EnHD single residue variants), experiments were performed in the forward direction to ensure adequate sampling of the native baseline. For proteins of higher stability (for example CHD single residue variants), experiments were performed in the reverse direction to ensure adequate sampling of the denatured baseline. For a subset of proteins, denaturation experiments were performed in both the forward and reverse directions to verify that the same conformational transitions were obtained for the two directions.

For each denaturation, titrant solution (protein in denaturant for the forward direction; protein in buffer for the reverse direction) was injected to a sample solution in a 1 cm path length cuvette (protein in buffer for the forward direction; protein in denaturant for the reverse direction). Samples were allowed to equilibrate for 5 minutes after each titrant injection. Following equilibration, the CD signal at 222 nm was averaged for 30 seconds at each GdnHCl concentration. Protein concentrations ranged from 2-12 μM. All titrations were performed at 20 °C and collected in triplicate for each variant.

Equilibrium folding free energies in the absence of denaturant (ΔG°_H2O_) and denaturant sensitivity coefficients (*m*-values) were determined using a two state linear extrapolation model (20). Because of the high C_m_ values of the most stable variants, determining ΔG°_H2O_ values requires a long extrapolation, which amplifies uncertainties that result from correlation between ΔG°_H2O_ and m-values. To minimize these uncertainties (21), we fit all unfolding curves for all variants to a global model (see Supplementary Materials, Fig S1 for a comparison of the global fits to local fits). In the global model, each unfolding curve was fit with local folded and unfolded baselines and a local ΔG°_H2O_ parameter, a single m-value parameter was shared among all curves and all variants. Prior to fitting, the CD signals were normalized such that the maximum signal intensity in each curve has a value of 1 and the minimum signal intensity has a value of 0 according to:

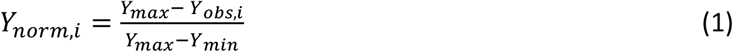

where Y_norm,i_ is the normalized signal of the *i*th data point in the curve, Y_max_ and Y_min_ are the maximum and minimum signal intensities respectively in the curve, and Y_obs,i_ is the signal intensity of the *ith* data point. For each variant, ΔG°_H2O_ values were averaged over the three replicate unfolding curves and uncertainties were determined as the standard errors of the mean. The difference in the stability of two variants was determined as the differences in folding free energies (ΔΔG°_H2O_) between the two variants. Uncertainties in ΔΔG °_H2O_ values were propagated from the uncertainties in ΔG°_H2O_ values according to:

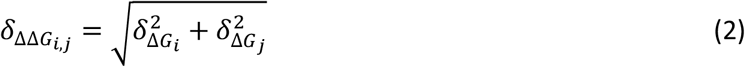

where 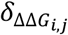 is the uncertainty in the ΔΔG°_H2O_ value for variant *i* and variant *j*, and 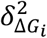 and 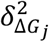 are the squared uncertainties in the ΔG°_H2O_ values of variant *i* and variant *j* respectively.

For EnHD, GdnHCl unfolding data is from a previous study (18). For CHD, we performed GdnHCl denaturations in the reverse direction to thoroughly sample the unfolded protein baseline and to maintain consistency with data collected here for CHD variants. The free energies and *m*-values for EnHD and CHD reported here are determined from the global model, and differ slightly from previously reported values (18).

### Residue solvent accessible surface area calculations

Residue-specific side chain solvent accessible surface areas (SASA) were calculated using the EnHD crystal structure (PDB: 1ENH) and were compared to the average solvent accessible surface area of the respective side chain in an ensemble of Gly-X-Gly tripeptides using GETAREA (22). Residues showing a relative side chain SASA less than 20% were classified as buried, residues showing a relative side chain SASA between 20-50% were classified as intermediate, and residues showing a relative side chain SASA greater than 50% were classified as surface positions. Residues D1, K2, K57, and K58 are not present in the EnHD crystal structure, but were classified as surface positions since they are the N- and C-terminal residues of the EnHD sequence. Two positions classified as buried contain charged resides in EnHD (E19 and K52). We reclassified these two positions with the intermediate group since both residues are right below the 20% side chain SASA (18.4% and 18.1% respectively, Table S2) and it is unlikely that the charged residues at these positions are completely buried.

### Analysis of feature overlaps

The overlap between features is determined as the number of substitutions shared between two features. The expected overlap (E) between features if the features are uncorrelated can be determined as the mean of a hypergeometric distribution (sampling substitutions without replacement) determined as:

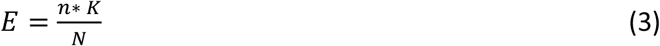

where n is the number of substitutions comprising feature 1, K is the number of substitutions comprising feature 2, and N is the total number of substitutions that are sampled (29 for all cases in the study).

## Results

### Substitutions by residue charge state

To explore whether consensus stabilization has electrostatic origins, we designed HD variants that combine substitutions that contribute to the charge differences between EnHD and CHD. In a study of seven consensus proteins, we found that consensus sequences often contain a higher percentage of charged residues than their naturally-occurring counterparts (16). This enhancement in charged residues is quite pronounced for CHD: CHD is made up of 50% charged residues, EnHD is made up of 40% charged residues (Fig 1A), and HDs on average are made up of ∼30% charged residues (16).

**Figure 1.**
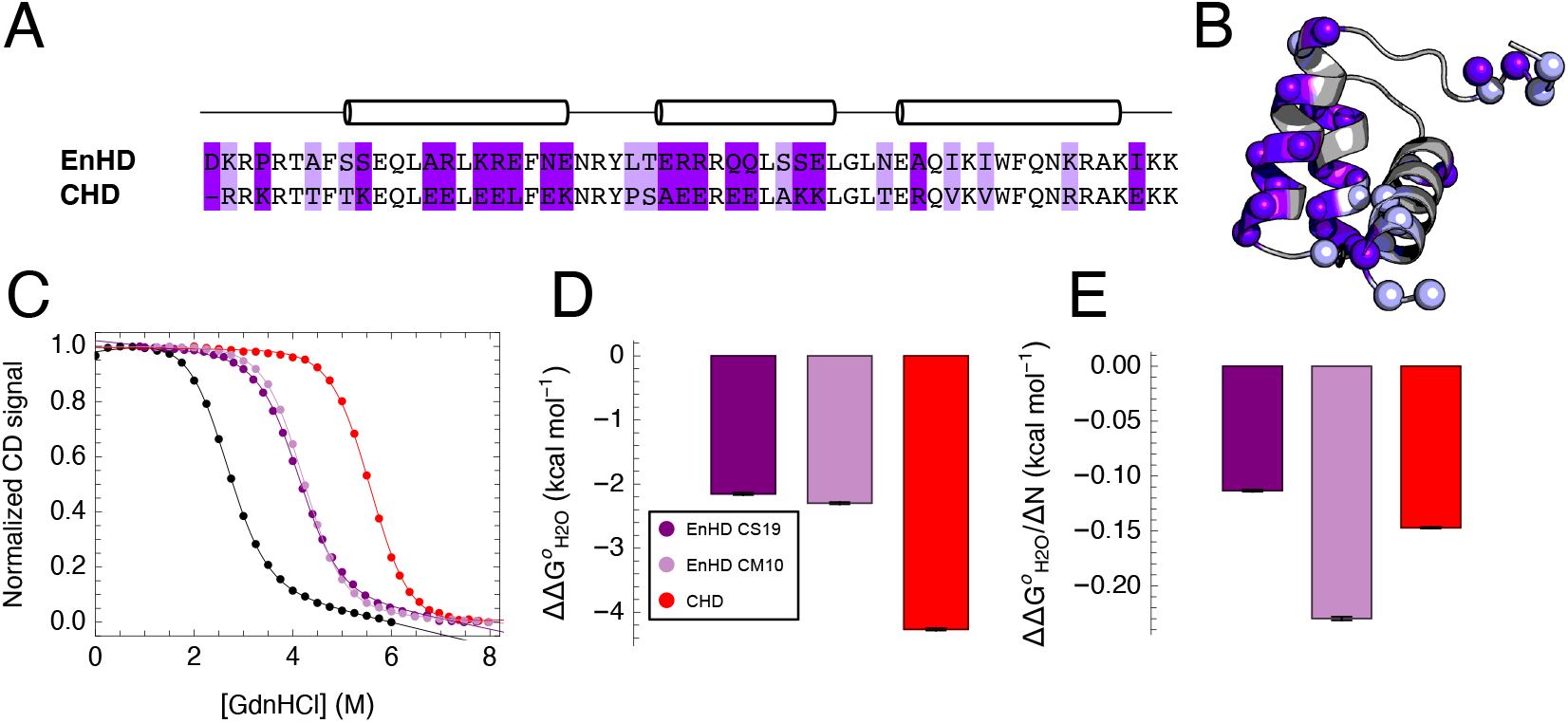
Consensus substitutions grouped by change in residue charge state. (A) Alignment of EnHD and CHD sequences. Sequence differences that change the charge state are shown in dark purple; differences that maintain the charge state are shown in lavender. Locations of α-helices are shown above the sequences. (B) Sequence differences mapped onto the EnHD structure (PDB 2JWT). Residues that differ between CHD and EnHD are shown with Cα atoms as spheres, and are colored as in panel A. (C) Representative GdnHCl-induced unfolding transitions for EnHD, EnHD CS19, EnHD CS10, and CHD (colored as in panel D). CD values are normalized to span from zero to one. Curves are two-state fits (Eq 1). Here and in all subsequent figures, data for EnHD (black) is from a previous study (18). (D) Effects on folding free energies relative to EnHD. Errors bars are uncertainties determined by Eq 2. (E) Effects on folding free energies relative to EnHD normalized for the number of substitutions (ΔN). Error bars are uncertainties from panel D divided by ΔN.

Of the 29 sequence differences between EnHD and CHD, 19 differences alter charge state (dark purple, Fig 1A, B). Nine of these 19 differences increase the net positive charge of CHD (negatively charged residues to a neutral or positively charged residue, or neutral residue to a positively charged residue). The other ten differences decrease the positive charge of CHD. Of the ten sequence differences that preserve the charge state, eight involve uncharged residues in both proteins; the other two substitute lysines with arginines (lavender, Fig 1A, B).

To examine the role of these charged residue differences in consensus stabilization, we designed a variant that introduces the consensus residues at the 19 positions that differ in charge state into EnHD, which we refer to as EnHD CS19 (for “**C**harge state **S**wapping”). We also made the complement to this set of substitutions, which differs from EnHD at the 10 positions that maintain the charge state, which we refer to as EnHD CM10 (for “**C**harge state **M**aintaining”).

EnHD CS19 and EnHD CM10 are both stabilized relative to EnHD (Fig 1C). Compared to EnHD, EnHD CS19 and EnHD CM10 have ΔΔG °_H2O_ values of -2.15 ± 0.02 and -2.30 ± 0.02 kcal mol^-1^, respectively (Fig 1D, Table S1). Thus, the charge-state maintaining and charge-state swapping consensus substitutions enhance stability to a roughly equal extent. However, neither the charge state swapping nor the charge state maintaining substitutions achieve the full 4.27 ± 0.02 kcal mol^-1^ increase in stability for all 29 consensus substitutions (Fig 1D). However, when stability increases are normalized for the number of substitutions in each set, the charge-state maintaining substitutions provide a greater stability enhancement per substitution than the charge-state swapping substitutions. The 10 charge-state maintaining substitutions provide -0.23 ± 0.01 kcal mol^-1^ per substitution, whereas the 19 charge state swapping substitutions provide an average of -0.11 ± 0.01 kcal mol^-1^ per substitution (Fig 1E).

### Substitutions by residue side chain burial

To determine the extent to which consensus stabilization results from substitution of buried versus solvent-exposed residues, we designed HD variants that combine substitutions at positions with similar degrees of side chain burial. The 29 sequence differences between EnHD and CHD occur at 19 surface positions, 7 intermediate positions, and 3 buried positions (Fig 2A, Fig 2B, Table S2).

**Figure 2.**
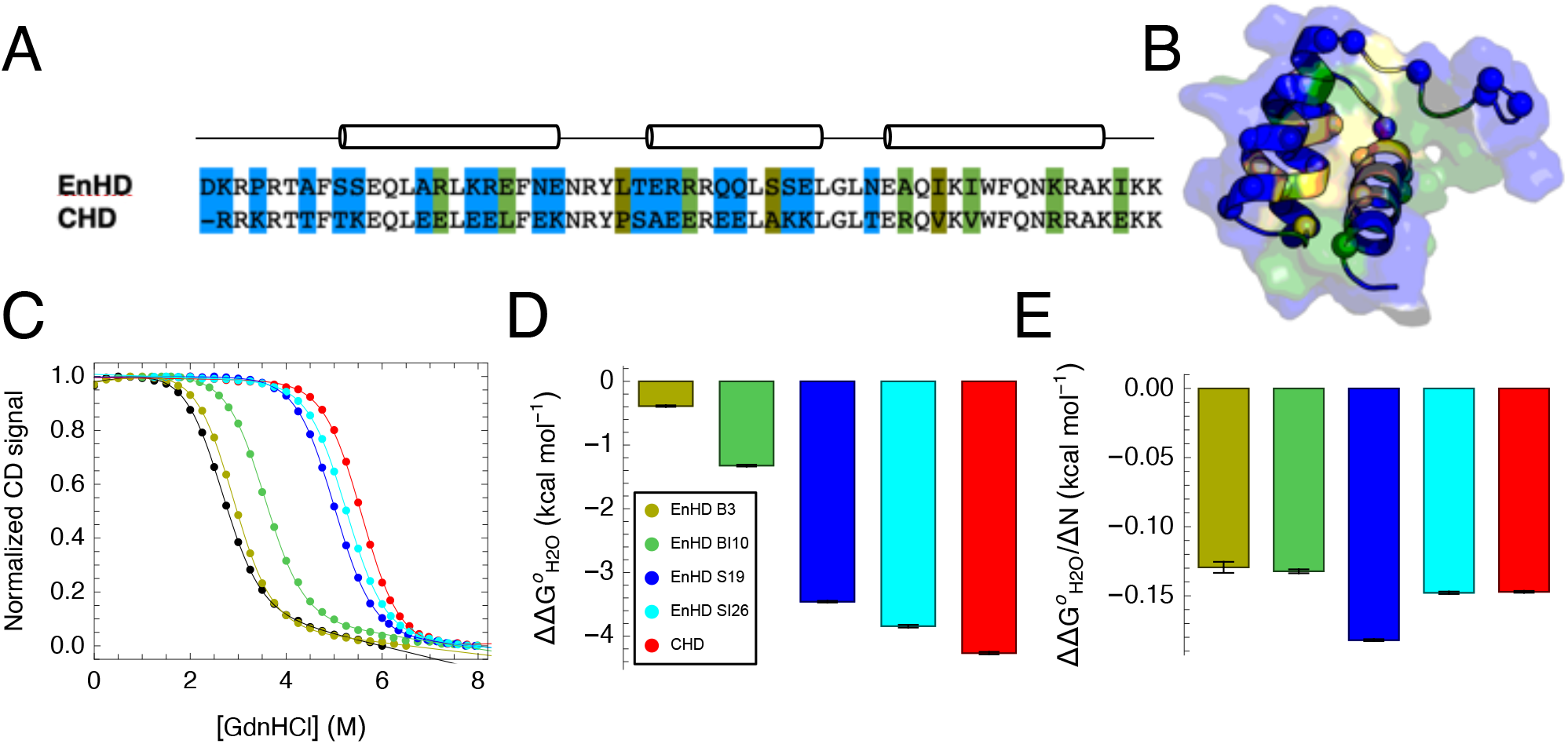
Consensus substitutions grouped by residue solvent accessibility in EnHD. (A) Alignment of EnHD and CHD sequences. Sequence differences at surface, intermediate, and buried sites are shown in blue, green, and yellow respectively. (B) Sequence differences mapped onto the EnHD structure. Residues that differ between EnHD and CHD are shown with Cα atoms as spheres and are colored as in panel A. (C) Representative GdnHCl-induced unfolding transitions for EnHD, EnHD B3, EnHD BI10, EnHD S19, EnHD SI26, and CHD. CD values are normalized to span from zero to one. Curves are two-state fits (Eq 1). Constructs are colored as in panel (D). (D) Effects on folding free energies relative to EnHD. Error bars are uncertainties determined by Eq 2. (E) Effects on folding free energies relative to EnHD normalized for the number of substitutions (ΔN). Error bars are uncertainties from panel D divided by ΔN.

We designed four variants that each introduce consensus residues with similar extents of burial into the EnHD background. EnHD B3 contains the three “**B**uried” consensus substitutions, EnHD BI 10 contains the 10 “**B**uried/**I**ntermediate” consensus substitutions, EnHD S19 contains the 19 “**S**urface” consensus substitutions, and EnHD SI26 contains the 26 “**S**urface/**I**ntermediate” substitutions (Fig 2C). Relative to EnHD, all four variants are stabilized. The buried and intermediate consensus substitutions show the smallest stability enhancement: the folding free energies of the EnHD B3 and EnHD BI10 variants are decreased by -0.39 ± 0.01 kcal mol^-1^ and -1.32 ± 0.01 kcal mol^-1^ compared to EnHD (Fig 2D, Table S1). In contrast, the surface consensus substitutions impart a greater stability enhancement: the folding free energy of the EnHD S19 variant is decreased by -3.46 ± 0.02 kcal mol^-1^ (Fig 2D, Table S1). The stabilizing contributions from the surface consensus substitutions are further underscored when stability increases are normalized for the number of substitutions in each set. Of the four variants that probe burial/accessibility, the surface consensus substitutions show the largest stability enhancement of 0.18 ± 0.01 kcal mol^-1^ per substitution, while the buried and buried/intermediately-exposed consensus substitutions each contribute 0.13 kcal mol^-1^ per substitution (Fig 2E).

### Substitutions by positional conservation

To determine the contributions of residue conservation to the consensus stability enhancement, we made variants that combine residues with different degrees of conservation. Conservation at a position *i* in an MSA was quantified using sequence information (SI_i_), calculated using the formula:

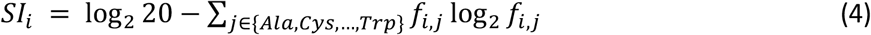

where *f*_*i,j*_ represents the frequency of residue *j* at position *i*. SI_i_ is the increase in entropy in going from the observed residue frequency distribution at position *i* (the sum term in the Eq 4) to a random distribution where all 20 residues have equal frequencies of 0.05. Positions with high SI are strongly conserved, whereas positions with low SI are weakly conserved.

We calculated SI for at each position of the homeodomain from an MSA composed of 4,571 sequences (see Supplementary Material). HD positions show a broad range of conservation, with values of SI ranging from 0.44 to 3.98, with a median value of 1.58 (Fig 3C). Positions with the same residues in EnHD and CHD (residues with black background in Fig 3A, black bars in Fig 3C) are on average more highly conserved (median SI of 2.12 bits) than positions that differ (residues with gray or colored backgrounds in Fig 3A, gray or colored bars in Fig 3C; median SI of 0.92 bits).

**Figure 3.**
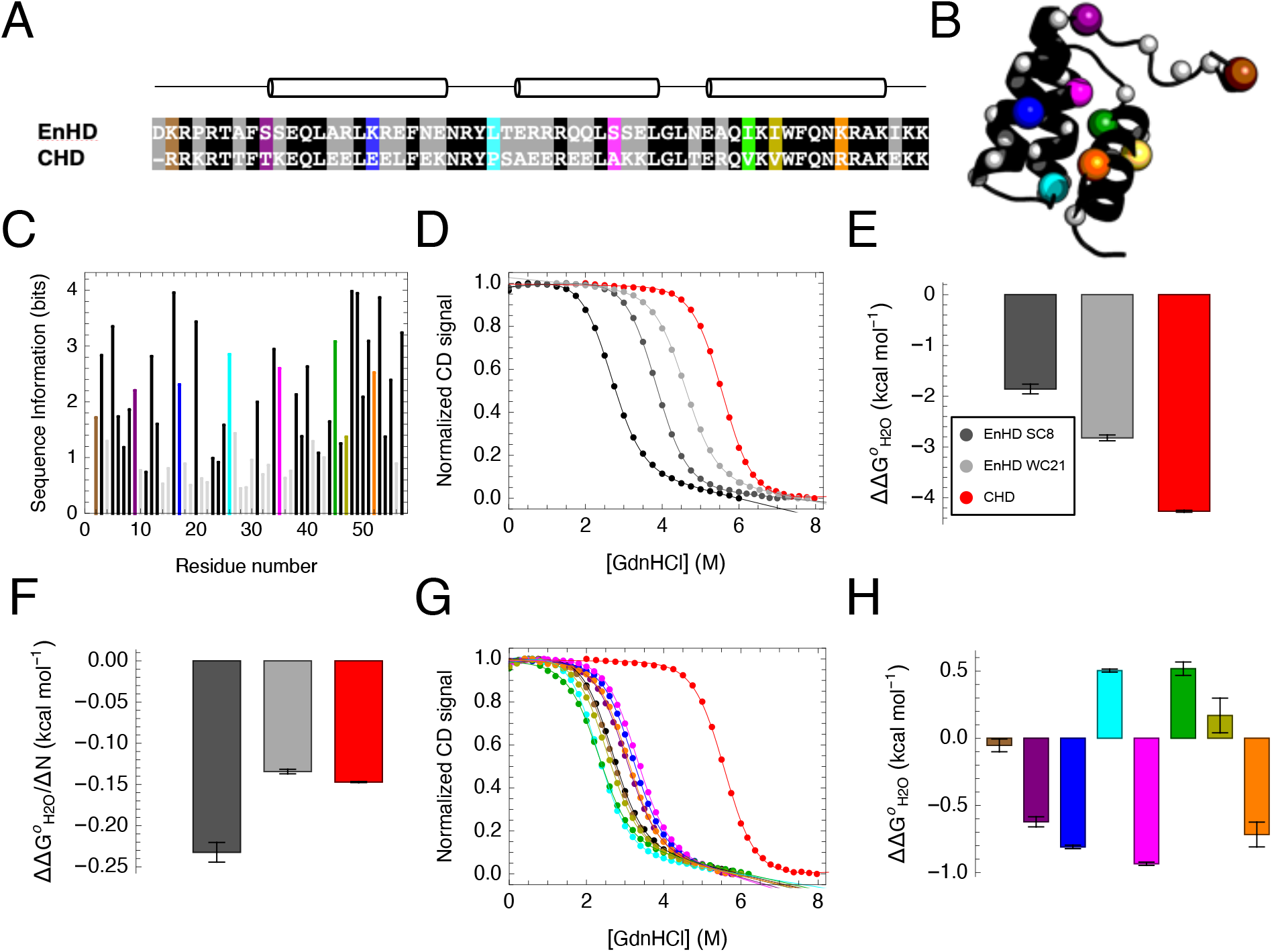
Consensus substitutions grouped by positional conservation. (A) Alignment of EnHD and CHD sequences. Residues with black backgrounds are identical in EnHD and CHD. Residues with grey background are the group of 21 differences at weakly conserved positions. Residues with other colors are the group of eight differences at strongly conserved positions. (B) Sequence differences mapped onto the EnHD structure. Cα atoms are shown as spheres and are colored as in panel A. (C) Sequence information (Eq 4) at each position from an alignment of 4,571 homeodomain sequences. Bars are colored as in panel A. (D) Representative GdnHCl-induced unfolding transitions for EnHD SC8 and EnHD WC21. (E) Effects of multiple residue variants on folding free energies relative to EnHD. (F) Effects on folding free energy changes relative to EnHD normalized for the number of substitutions (ΔN). Red and black transitions (panels D, G) are for EnHD and CHD. (G) Representative GdnHCl-induced unfolding transitions for EnHD single residue variants (colored as in panel A). (H) Effects of single residue substitutions in EnHD on folding free energies. Error bars in panels E and H are uncertainties determined by Eq 2. Error bars in panel F are uncertainties from panel E divided by ΔN.

However, not all positions that differ between EnHD and CHD have low SI values. Seven such positions have SI values greater than the median sequence information across all positions. To the extent that effects on stability correlate with residue frequencies, consensus substitutions at positions with high SI values may be expected to show the greatest increases in stability.

To determine the contributions to stability from residues at positions of high conservation, we designed a variant that combines consensus substitutions at eight of the most strongly conserved positions into the EnHD background, which we refer to as EnHD SC8 (for eight **S**trongly **C**onserved). We also made the complementary variant which introduces the consensus substitutions at the remaining 21 “**W**eakly **C**onserved” positions, which we refer to as EnHD WC21.

EnHD SC8 and EnHD WC21 both show unfolding transitions intermediate between EnHD and CHD (Fig 3D). EnHD SC8 and EnHD WC21 show decreases in folding free energies relative to EnHD of -1.86 ± 0.10 kcal mol^-1^ and -2.82 ± 0.06 kcal mol^-1^ respectively, indicating that both sets of substitutions are stabilizing, with the weakly conserved substitutions being slightly more stabilizing (Fig 3E, Table S1). When these free energy changes are normalized for the number of substitutions, the strongly conserved substitutions decrease the folding free energy by -0.23 ± 0.01 kcal mol^-1^ per substitution, whereas the weakly conserved substitutions decrease the folding free energy decrement by -0.13 ± 0.01 kcal mol^-1^ per substitution (Fig 3F). Although it is not unexpected that consensus substitutions at positions of high conservation contribute more to stability on average than at positions of low conservation, the larger number of sequence differences at weakly conserved positions means that a considerable stabilization is provided by consensus substitution at weakly conserved positions.

To further explore the relationship between conservation and stability, we measured the stabilities of individual substitutions from the eight strongly conserved positions in the EnHD background (Fig 3G, Table S1). Four of the eight strongly conserved substitutions are stabilizing, three are destabilizing (though the destabilizing effect of I47V is quite small), and one has no effect on stability. On the whole, changes in folding free energies for these eight conserved point-substitutions are small: ΔΔG °_H2O_ values from EnHD range from -0.93 ± 0.01 kcal mol^-1^ to +0.52 ± 0.05 kcal mol^-1^ with a mean value of -0.24 kcal mol^-1^ (Fig 3H, Table S1), clearly demonstrating that the consensus stability enhancement (−4.27 ± 0.02 kcal mol^-1^) is not gained through a small number of substitutions at conserved positions.

### Nonadditive effects on stability of consensus substitutions

The full consensus HD has 29 sequence differences from *D. melanogaster* Engrailed HD, which is half of the residues in the protein. Thus, although CHD and EnHD have similar tertiary structures (18), they are quite different in sequence. Many of these differences involve residues that are in direct contact (Table S3). Residues in direct contact often make non-additive contributions to protein stability (23). To determine whether non-additivity contributes to consensus stability enhancement, we made individual residue substitutions at the eight strongly conserved sites in the CHD background (Fig 4A), so that we could compare the stability changes to those measured in the EnHD background (Fig 3G). For example, we made the E17K substitution in the CHD background to compare to the K17E substitution made in the EnHD background, thereby allowing us to test the effects of the consensus versus non-consensus background on stabilization from individual consensus substitutions.

**Figure 4.**
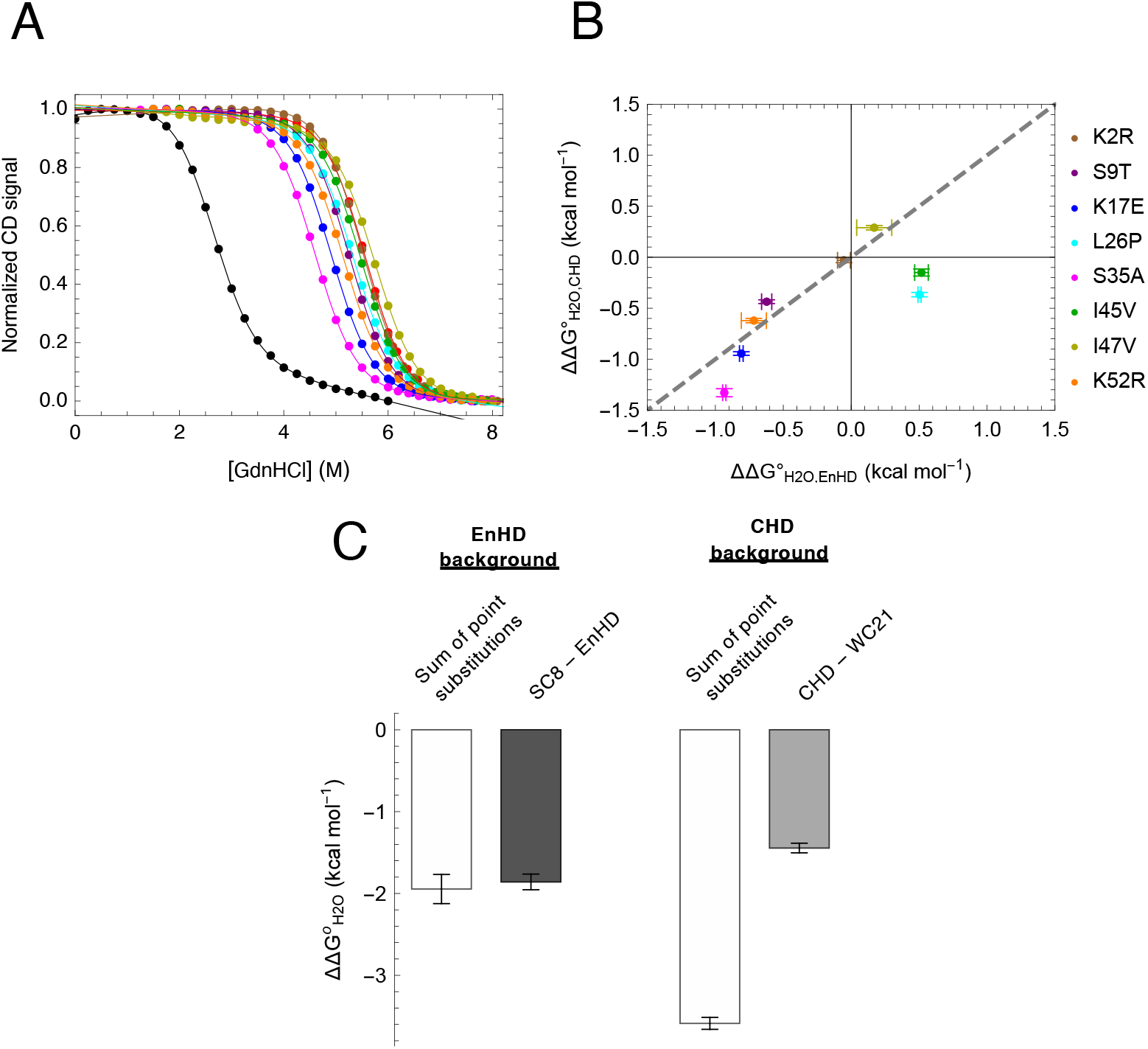
Non-additive effects of consensus substitutions on stability. (A) Representative GdnHCl-induced unfolding transitions for CHD single residue variants along with EnHD (black) and CHD (red). Unfolding transitions are colored as in panel B. (B) Correlation of effects on folding free energies of single residue substitutions in EnHD background (x-axis) and CHD background (y-axis). Dashed gray line shows the y=x relationship. ΔΔG°_H2O_ values are determined from a global fit with a common m-value (see text). Error bars are uncertainties determined by Eq 2. (C) Additivities of the folding free energies of the eight highly conserved residue substitutions in the EnHD and CHD backgrounds. White bars are the sum of ΔΔG°_H2O_ values for the eight single residue variants in the EnHD (left) and CHD (right) backgrounds. Error bars are uncertainties propagated from ΔΔG°_H2O_ values of the single residue variants. Gray bars are ΔΔG°_H2O_ values when all eight substitutions are made simultaneously in the EnHD and CHD backgrounds; error bars are uncertainties determined by Eq 2.

To compare the effects of substitution on the folding free energies in the EnHD and CHD backgrounds, we determined ΔΔG °_H2O_ values in each background for substitution *toward* the consensus sequence by:

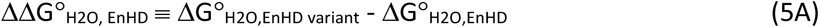

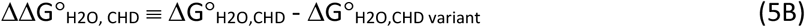

where ΔΔG°_H2O, EnHD_ is the effect of the substitution (toward consensus) in the EnHD background and ΔΔG°_H2O, CHD_ is the effect of the substitution (away from consensus) in the CHD background. With these definitions, substitutions that introduce consensus residues that are stabilizing have negative ΔΔG°_H2O_.

In the CHD background, six substitutions toward consensus residue have negative ΔΔG°_H2O_ values (although the stabilizing effect of the I45V substitution is quite small, -0.15 ± 0.04 kcal mol^-1^), whereas only four are stabilizing in the EnHD background (Figs 4A-B). Thus, two consensus substitutions (L26P and I45V) that are destabilizing in the EnHD background become destabilizing in the CHD background. Furthermore, whereas the A35S is stabilizing by 0.93 ± 0.01 kcal mol^-1^ in the EnHD background, it is stabilizing by 1.33 ± 0.01 kcal mol^-1^ in the CHD background. Thus, the consensus background enhances the stabilizing effects of these three consensus substitutions. This non-additivity is not the result of using a shared m-value in fits using a global model: stabilization in the consensus background is even larger when determined using the free energies from the local fits (Fig S2).

Non-additive effects of consensus residue substitutions can also be seen by comparing stability enhancement of the combined SC8 substitution with the sum from single-residue substitutions in the EnHD and CHD backgrounds. The sum of ΔΔG°_H2O_ values for the eight single residue substitutions in the EnHD background (−1.95 ± 0.08 kcal mol^-1^) is nearly the same as the ΔΔG°_H2O_ value for EnHD SC8 relative to EnHD (−1.86 ± 0.10 kcal mol^-1^), closely approximating additivity (Fig 4C). In contrast, the sum of ΔΔG°_H2O_ values for the eight single residue substitutions in the CHD background (−3.59 ± 0.07 kcal mol^-1^) is larger than the ΔΔG°_H2O_ value for CHD relative to EnHD WC21 (−1.44 ± 0.06 kcal mol^-1^; Fig 4C), demonstrating clear non-additivity. Thus, on average, consensus substitutions are more stabilizing in the full consensus background than they are in a protein that is eight substitutions away from the consensus.

## Discussion

The large enhancement in stability of CHD relative to EnHD must arise from the 29 residue differences between the two proteins. However, it is unclear how this stability enhancement is partitioned among the 29 substitutions. Do a small subset of substitutions contribute the bulk of the stability enhancement? Are there particular types of substitutions that contribute the bulk of the stability enhancement? Are these contributions additive? By introducing sets of residue substitutions with shared sequence and/or structural features, we can assess and rank the contributions of these features to the consensus stability enhancement.

### Overlap between groups of substitutions

When comparing changes in folding free energies for the different sets of substitutions, the degree of overlap between different groups needs to be considered. Because complementary groups of substitutions (for example, charge swapping and charge maintaining) together comprise all 29 sequence differences between EnHD and CHD, overlap between substitution groups that probe different features is unavoidable. The strongest overlap is between the charge-swapping and weakly conserved substitutions: 18 of the 19 charge swapping substitutions are also in the weakly conserved group of 21 substitutions (Fig 5A). Indeed, the CS19 and WC21 variants differ by only four substitutions (Fig 5A, Table S4). This is partly a result of the large number of substitutions in each group (19 and 21); we would expect an overlap of about 14 substitutions if these two features sorted independently (see Methods).

**Figure 5.**
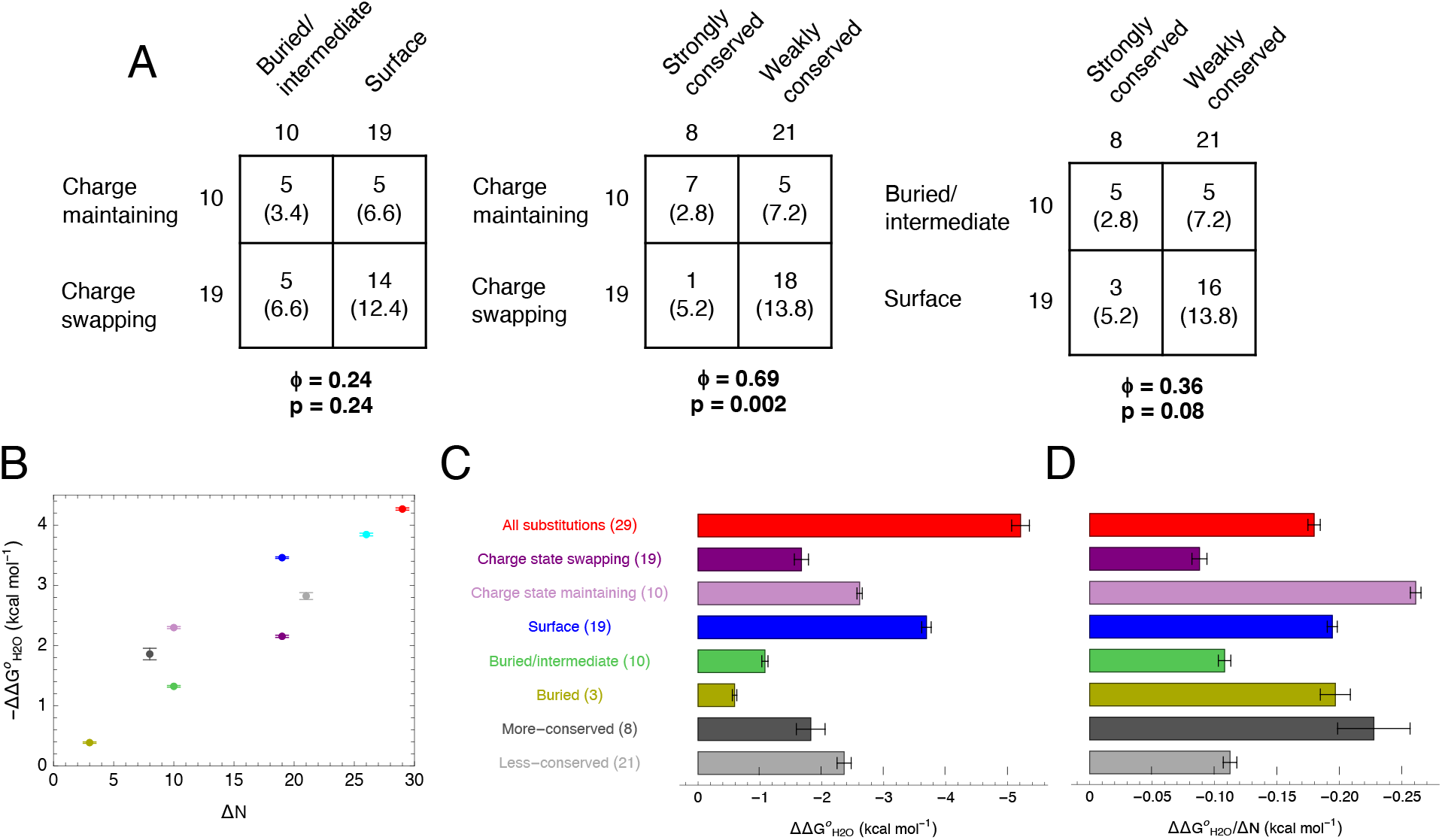
Stability contributions of substitutions of residues in different sequence/structure classes. (A) Contingency tables for the overlap of the residue substitutions for each feature. The top number indicates the number of substitutions in shared by the two features. The bottom number in parentheses indicates the expected shared number of substitutions if the features were uncorrelated determined by Eq 3. The Matthews correlation coefficient (ϕ) between features is given below each table. The p-value of feature correlation is determined by Fisher’s exact test. (B) Correlation of effects on folding free energies with the number of substitutions encompassed for each feature. (C) Total effects and (D) per-substitution effects on stability from consensus substitutions into EnHD for each feature. Error bars in panels B and C are uncertainties determined by Eq 2. Error bars in panel D are uncertainties from panel C divided by the number of substitutions for each variant.

Likewise, there is considerable overlap between strongly conserved and charge-maintaining substitutions: 7 out of 8 strongly conserved substitutions are charge maintaining, whereas a four residue overlap would be expected from independent assortment. Together, these overlaps produce a Matthews correlation coefficient (ϕ) of 0.68 for the charge versus conservation groups.

For variants that probe solvent exposure (BI10 and S19) and conservation (SC8 and WC21), overlaps are closer to what would be expected from random assortment (Fig 5A). Correspondingly, Matthews correlation coefficients among these two groups are lower (0.36 and 0.24, respectively). The largest overlap among these groups is that 16 of the 19 surface substitutions are in the weakly conserved weakly conserved group of 21 substitutions. Though this overlap is only two residues in excess of what would be predicted from random assortment, it does mean that these two groups are largely reporting on the same substitutions; thus, differences in stability between S19 and WC21 come from a small group of substitutions that probe one but not the other feature.

### Comparison of stabilizing effects of different sequence and structural features

Across the three features we examined—residue charge state, residue burial, and conservation—all sets of consensus substitutions were found to be stabilizing. Thus, stabilizing effects from consensus substitutions do not appear to be limited to a specific feature. For all variants we tested, we see a strong positive correlation between the extent of stabilization and the number of substitutions made (Fig 5B). This suggests that consensus stabilization may arise from small, incremental effects from substitutions regardless of features, and that a simple way to gain additional stability may be to simply include more consensus substitutions. However, this correlation is not absolute. For example, although both the charge state swapping substitutions and the surface substitutions each contain 19 substitutions, the surface substitutions account for 1.3 kcal mol^-1^ more stability than the charge state swapping substitutions (Fig 5C).

In terms of per-residue effects, the largest stabilization is seen for the charge-maintaining and strongly conserved substitutions (−0.23 kcal mol^-1^ residue^-1^ for each group; Fig 5D), although these groups are strongly overlapping. This indicates that if a high stability increment is to be achieved from a limited number of consensus substitutions, targeting conserved sites that maintain charge seems a good bet. In terms of total effects, the largest stabilization is seen for the surface (S19) and weakly-conserved (WC21) groups (ΔΔG°_H2O_ = -3.46 and -2.82 kcal mol^-1^, respectively), demonstrating that a large portion of consensus stabilization comes from what seems to be unlikely sources—residues that minimally constrained, both structurally and evolutionarily. The large overall contribution from the weakly conserved substitutions (Fig 5C) indicates that even modest sequence biases reflect stability, and is consistent with findings from Swint-Kruse and coworkers that highlight the importance of weakly-conserved surface residues for protein function (24, 25).

Comparison of complementary sets of substitutions within a single feature provides a measure of stability that is, by definition, free from overlap. Per residue, greater stabilization is afforded from consensus substitutions at strongly conserved sites than from substitutions at weakly conserved sites (ΔΔG°_H2O_ /ΔN = -0.23 versus -0.13 kcal mol^-1^ residue^-1^), from surface consensus substitutions rather than buried and intermediately-exposed substitutions (ΔΔG°_H2O_ /ΔN = -0.18 versus -0.13 kcal mol^-1^ residue^-1^), and from charge-maintaining consensus substitutions than from charge-swapping substitutions (ΔΔG°_H2O_ /ΔN = -0.23 versus -0.11 kcal mol^-1^ residue^-1^). This latter observation suggests that even though consensus proteins are enriched in ionizable residues (16), consensus stabilization is not primarily the result of stabilizing charge-charge interactions.

### Contribution of non-additivity to consensus stabilization

A comparison of identical consensus substitutions in the EnHD and CHD backgrounds reveals favorable couplings among residues in the consensus background compared to the EnHD background. Whereas five out of eight single-residue substitutions (K2R, S9T, K17E, I47V, and K52R) show similar effects on stability in both backgrounds, three substitutions (L26P, S35A, and I45V) show a larger stabilizing effect when introduced into the CHD background than the EnHD background (Fig 4B).

Moreover, two of these background-dependent substitutions (L26P and I45V) are destabilizing in the EnHD background but become stabilizing (albeit marginally so) in the CHD background (Fig 4B). This indicates mutual stabilization between the each of the L26P, S35A, and I45V substitutions with the other 28 substitutions that define the consensus HD background.

Formally, the background dependence of the ΔΔG°_H2O_ values for the L26P, S35A, and I45V substitutions represents the coupling of these residues to the entire EnHD and CHD backgrounds, which differ by the remaining 28 mismatching positions. We cannot identify the specific interactions that give rise to this background dependence, though non-additive effects of substitutions typically occur between residues in direct contact (in some cases, longer range interactions can result from electrostatic effects or structural perturbations caused by the substitutions) (23). Among the eight individual substitutions we tested, the residues at the three positions showing non-additivity make either two or three side chain-side chain contacts with other residues in the EnHD structure (Table S3). The residues at the other five positions make at most one side chain-side chain contact. Furthermore, the residues at the three positions showing non-additivity make contacts with one or both of the other positions showing non-additivity. Interestingly, all three background-dependent substitutions are the three buried substitutions in the protein core (Figs 2A, 4B). This suggests a potential mechanism underlying consensus protein stability enhancement involving favorable couplings among moderately conserved groups of residues in the protein core that are absent, on average, in the core of a natural proteins.

Three previous studies have explored non-additivity in consensus stabilization. Fersht and coworkers found that p53 DNA-binding domain variants containing five different combinations of the same four consensus substitutions showed additive effects (17). However, these four substitutions chosen were at positions distant in structure (ranging from 11-24 Å away) and thus additive effects among the substitutions may have been expected (23). Likewise, in a separate study Fersht and coworkers found that consensus substitutions in two variants containing six consensus substitutions in *E. coli* GroEL minichaperones were additive; notably, some of these substitutions were close in space (26). In contrast, Magliery and coworkers found that consensus substitutions in a triosephosphate isomerase (TIM) were non-additive, albeit in the opposite direction we have observed here: combining 13 consensus substitutions (plus an additional non-consensus substitution) that were all individually stabilizing resulted a variant that was slightly destabilized relative to the wild type TIM, suggesting that consensus substitutions synergistically destabilize one another (27).

In addition to the coupling of individual substitutions to the CHD versus EnHD background, we see clear non-additivity among the eight consensus substitutions in the CHD background. When the eight strongly conserved consensus substitutions are introduced individually into in the full consensus background, the sum of the stability enhancements exceeds that obtained when all eight substitutions are made simultaneously (comparing CHD to WC21; Fig 4C). In this format, the single substitutions in the consensus background can each be thought of as a “last substitution”, where all other non-consensus residues have been replaced with consensus residues. This observation indicates that the eight strongly conserved substitutions mutually stabilize one another in the consensus background. The observation that the same eight substitutions appear to be additive in the EnHD environment (comparing the sum of the single substitutions SC8 minus EnHD, Fig 4C) indicates that the synergy among the eight strongly conserved residues also requires consensus residues at the 21 positions of lower conservation, indicating that non-additivity includes “higher-order” couplings involving three or more residues.

Furthermore, the observation that individual consensus substitutions are more stabilizing in the consensus background than in the EnHD background suggests that making individual consensus substitutions in a naturally occurring sequence may not provide much stabilization, as has been seen in a number of studies (15, 17, 26, 28).

Superficially, the finding of stabilizing energetic couplings between residues in a consensus protein is somewhat surprising, since the consensus design method selects residues at each position without regard for sequence at any other position. How are these energetic couplings are captured in a consensus sequence? If stabilizing energetic couplings are selected by evolution, the energetic couplings may give rise to positive sequence covariance between favorably coupled residues. Although covariance information is not explicitly used in consensus design, positive covariance should enhance frequencies for both covariant residues. Thus, positive covariances may be implicitly encoded within the conservation of residues at a single position, and consensus sequences may capture these covariant residues. However, the extent to which sequence covariance reflects stabilizing interactions remains an open question. Ranganathan and co-workers have demonstrated that including residues that co-evolve is necessary for proteins to adopt their native folds (29). However, the results from Magliery and co-workers described above demonstrate that excluding residues that co-evolve improves the identification of stabilizing substitutions (27). The role of sequence covariance in stabilizing (or destabilizing) consensus-based sequences is a topic worthy of further study.

## Supporting information

Supplemental material for Sternke et al.

## Author contributions

M. S., K. W. T., and D. B. designed the research. M. S. and K. W. T. performed the research. M. S., K. W. T., and D. B. analyzed data. M. S., K. W. T., and D. B. wrote the manuscript.

## Acknowledgements

The authors would like to thank the Johns Hopkins University Center for Molecular Biophysics for providing facilities, instrumentation, and resources. This work was supported by NIH/NIGMS research grant R01 GM068462 to D.B., NIH/NIGMS training grant T32 GM008403 for M.S., and NIH/NIGMS fellowship F31 GM128295 to M.S.

## References

1. Taverna, D.M., and R.A. Goldstein. 2002. Why are proteins marginally stable? Proteins. 46:105– 109.

2. Zeldovich, K.B., P. Chen, and E.I. Shakhnovich. 2007. Protein stability imposes limits on organism complexity and speed of molecular evolution. Proc. Natl. Acad. Sci. U. S. A. 104:16152–16157.

3. Razvi, A., and J.M. Scholtz. 2006. Lessons in stability from thermophilic proteins. Protein Sci. 15:1569–1578.

4. Koga, N., R. Tatsumi-Koga, G. Liu, R. Xiao, T.B. Acton, G.T. Montelione, and D. Baker. 2012. Principles for designing ideal protein structures. Nature. 491:222–227.

5. Itzhaki, L.S., D.E. Otzen, and A.R. Fersht. 1995. The Structure of the Transition State for Folding of Chymotrypsin Inhibitor 2 Analysed by Protein Engineering Methods: Evidence for a Nucleation-condensation Mechanism for Protein Folding. J. Mol. Biol. 254:260–288.

6. Nisthal, A., C.Y. Wang, M.L. Ary, and S.L. Mayo. 2019. Protein stability engineering insights revealed by domain-wide comprehensive mutagenesis. Proc. Natl. Acad. Sci. 116:16367–16377.

7. Geiger-Schuller, K., K. Sforza, M. Yuhas, F. Parmeggiani, D. Baker, and D. Barrick. 2018. Extreme stability in de novo-designed repeat arrays is determined by unusually stable short-range interactions. Proc. Natl. Acad. Sci. 115:7539–7544.

8. Mravic, M., J.L. Thomaston, M. Tucker, P.E. Solomon, L. Liu, and W.F. DeGrado. 2019. Packing of apolar side chains enables accurate design of highly stable membrane proteins. Science. 363:1418– 1423.

9. Huang, P.-S., G. Oberdorfer, C. Xu, X.Y. Pei, B.L. Nannenga, J.M. Rogers, F. DiMaio, T. Gonen, B. Luisi, and D. Baker. 2014. High thermodynamic stability of parametrically designed helical bundles. Science. 346:481–485.

10. Giver, L., A. Gershenson, P.O. Freskgard, and F.H. Arnold. 1998. Directed evolution of a thermostable esterase. Proc. Natl. Acad. Sci. U. S. A. 95:12809–12813.

11. Arnold, F.H. 2015. The nature of chemical innovation: new enzymes by evolution*. Q. Rev. Biophys. 48:404–410.

12. Broom, A., K. Trainor, Z. Jacobi, and E.M. Meiering. 2020. Computational Modeling of Protein Stability: Quantitative Analysis Reveals Solutions to Pervasive Problems. Structure. 28:717-726.e3.

13. Porebski, B.T., and A.M. Buckle. 2016. Consensus protein design. Protein Eng. Des. Sel. PEDS. 29:245–251.

14. Akanuma, S., Y. Nakajima, S. -i. Yokobori, M. Kimura, N. Nemoto, T. Mase, K. -i. Miyazono, M. Tanokura, and A. Yamagishi. 2013. Experimental evidence for the thermophilicity of ancestral life. Proc. Natl. Acad. Sci. 110:11067–11072.

15. Sternke, M., K.W. Tripp, and D. Barrick. 2020. The use of consensus sequence information to engineer stability and activity in proteins. In: Tawfik DS, editor. Methods in Enzymology. Academic Press. pp. 149–179.

16. Sternke, M., K.W. Tripp, and D. Barrick. 2019. Consensus sequence design as a general strategy to create hyperstable, biologically active proteins. Proc. Natl. Acad. Sci. 116:11275–11284.

17. Nikolova, P.V., J. Henckel, D.P. Lane, and A.R. Fersht. 1998. Semirational design of active tumor suppressor p53 DNA binding domain with enhanced stability. Proc. Natl. Acad. Sci. U. S. A. 95:14675–14680.

18. Tripp, K.W., M. Sternke, A. Majumdar, and D. Barrick. 2017. Creating a Homeodomain with High Stability and DNA Binding Affinity by Sequence Averaging. J. Am. Chem. Soc. 139:5051–5060.

19. Nozaki, Y. 1972. The preparation of guanidine hydrochloride. Methods Enzymol. 26:43–50.

20. Street, T.O., N. Courtemanche, and D. Barrick. 2008. Protein Folding and Stability Using Denaturants. In: Methods in Cell Biology. Academic Press. pp. 295–325.

21. Marold, J.D., K. Sforza, K. Geiger-Schuller, T. Aksel, S. Klein, M. Petersen, E. Poliakova-Georgantas, and D. Barrick. 2020. A collection of programs for one-dimensional Ising analysis of linear repeat proteins with point substitutions. bioRxiv. 2020.06.27.175224.

22. Fraczkiewicz Robert and Braun Werner. 1998. Exact and efficient analytical calculation of the accessible surface areas and their gradients for macromolecules. J. Comput. Chem. 19:319–333.

23. Wells, J.A. 1990. Additivity of mutational effects in proteins. Biochemistry. 29:8509–8517.

24. Fenton, A.W., B.M. Page, A. Spellman-Kruse, B. Hagenbuch, and L. Swint-Kruse. 2020. Rheostat positions: A new classification of protein positions relevant to pharmacogenomics. Med. Chem. Res. 29:1133–1146.

25. Meinhardt, S., M.W.J. Manley, D.J. Parente, and L. Swint-Kruse. 2013. Rheostats and toggle switches for modulating protein function. PloS One. 8:e83502.

26. Wang, Q., A.M. Buckle, N.W. Foster, C.M. Johnson, and A.R. Fersht. 1999. Design of highly stable functional GroEL minichaperones. Protein Sci. 8:2186–2193.

27. Sullivan, B.J., T. Nguyen, V. Durani, D. Mathur, S. Rojas, M. Thomas, T. Syu, and T.J. Magliery. 2012. Stabilizing Proteins from Sequence Statistics: The Interplay of Conservation and Correlation in Triosephosphate Isomerase Stability. J. Mol. Biol. 420:384–399.

28. Di Nardo, A.A., S.M. Larson, and A.R. Davidson. 2003. The Relationship Between Conservation, Thermodynamic Stability, and Function in the SH3 Domain Hydrophobic Core. J. Mol. Biol. 333:641–655.

29. Socolich, M., S.W. Lockless, W.P. Russ, H. Lee, K.H. Gardner, and R. Ranganathan. 2005. Evolutionary information for specifying a protein fold. Nature. 437:512–518.

