## Supplemental material for Sternke et al. for "Surface residues and non-additive interactions stabilize a consensus homeodomain protein"

**Supplementary text**

**Comparison of global and local fits**

Comparing $\Delta\Delta G°_{H2O}$ values among EnHD, CHD and all variants is complicated due to the long guanidine extrapolations required for the variants with high stabilities. The range of m-values from local fits of the unfolding transition are all within 0.3 kcal $mol^{-1}$ $M^{-1}$ (Table S1), however these small variations have significant effects on $\Delta G°_{H2O}$ values owing to the long extrapolation from the unfolding midpoints (~4.5 to 6 M GdnHCl) for the most-stable of the variants. Thus, interpreting changes in stability based on changes in $\Delta G°_{H2O}$ values may lead to inaccurate $\Delta\Delta G°_{H2O}$ values, especially for single-residue variants in the consensus background.

To minimize the effects that *m*-value differences have on extrapolated folding free energies, we globally fit the unfolding transitions for EnHD, CHD, and all variants to a model containing a single global *m*-value, but local $\Delta G°_{H2O}$ and baseline parameters for each unfolding transition. This restrained model fits reasonably well to the data; residuals are randomly dispersed and are only slightly larger than the residuals from the local fits (Fig S1). Corrected for the difference in the number of degrees of freedom, squared residuals for the global m-value fit are increased by a factor 1.62 compared to combined residuals from the local fits (Fig S1). Folding free energies from the two models are quite similar, with a root mean square deviation of 0.37 kcal $mol^{-1}$ over all proteins; this is between 4-10% of the magnitude of the measured folding free energies. Moreover, the trends in the stabilizing effects for all variants, including the single residue substitutions and larger substitution sets, all remain consistent between the two models (Table S1).

### Supplementary Figures and Tables

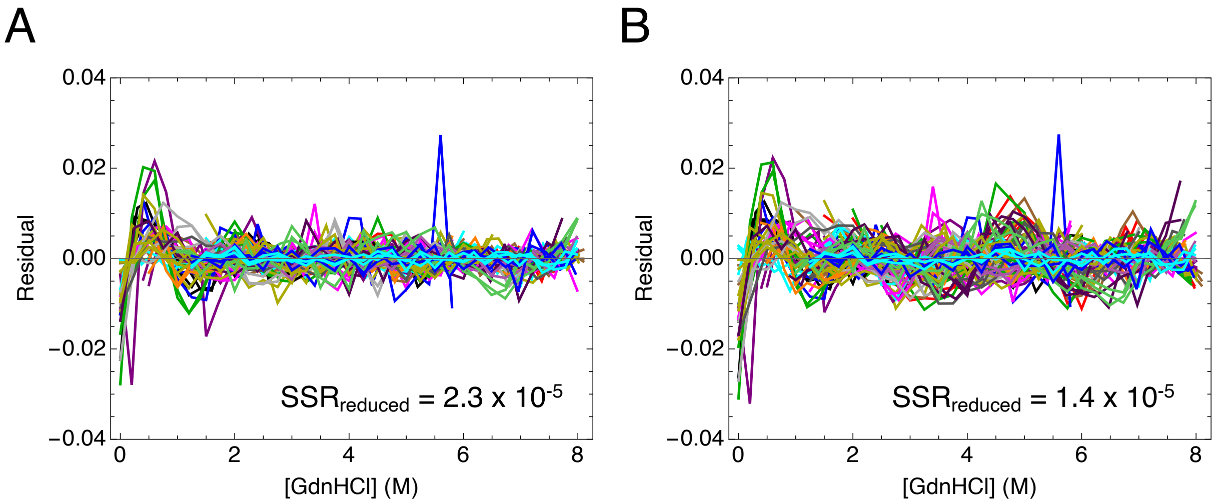

**Figure S1. Comparing global and local fits of unfolding transitions.** Residuals for the fits of all unfolding transitions for all proteins using (A) the global model and (B) the local model. Unfolding transitions and fits of the global model are shown in Figs 1-4. Residuals are colored as in Figs 1-4. $SSR_{reduced}$: sum of squared of residuals per degree of freedom.

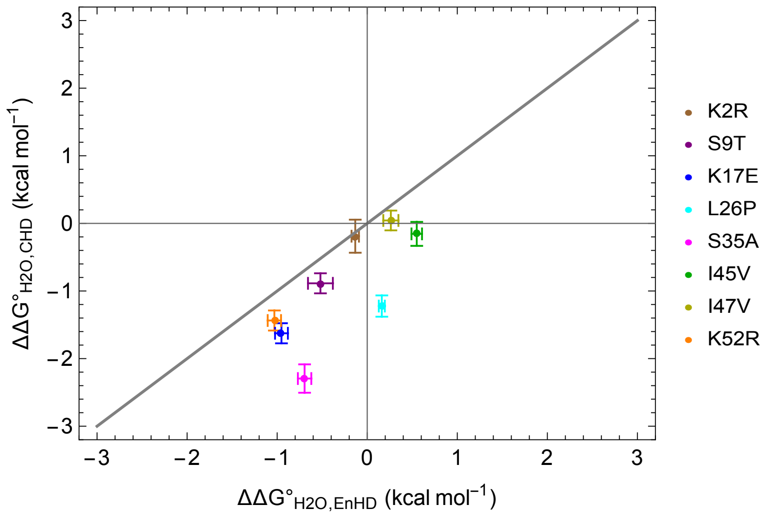

**Figure S2. Background effects of residue substitutions from local fits.** Correlation of effects on folding free energies of single residue substitutions toward the consensus in EnHD background and CHD background. Diagonal line represents a one-to-one (y=x) relationship. Folding free energies are determined from local fits of all proteins (data in Table S1). Errors in $\Delta\Delta G^{\circ}_{H2O}$ are determined by Eq 2.

**Table S1. Folding thermodynamics of EnHD and CHD variants.**

| Protein | $\Delta G^{\circ}_{H2O}$[a] (kcal mol$^{-1}$) | $C_m$[a] (M) | $\Delta\Delta G^{\circ}_{H2O}$[b] (kcal mol$^{-1}$) | $\Delta N$[c] | $\Delta\Delta G^{\circ}_{H2O}/\Delta N$[d] (kcal mol$^{-1}$) | $\Delta G^{\circ}_{H2O,local}$[e] (kcal mol$^{-1}$) | *m*-value (kcal mol$^{-1}$ M$^{-1}$)[e] |
|---|---|---|---|---|---|---|---|
| EnHD | -3.76 ± 0.01 | 2.60 ± 0.01 | --- | --- | --- | -3.62 ± 0.03 | 1.39 ± 0.01 |
| CHD | -8.03 ± 0.02 | 5.56 ± 0.02 | -4.27 ± 0.02 | 29 | -0.15 ± 0.01 | -8.83 ± 0.14 | 1.59 ± 0.02 |
| EnHD CS19 | -5.91 ± 0.02 | 4.10 ± 0.01 | -2.15 ± 0.02 | 19 | -0.11 ± 0.01 | -5.29 ± 0.11 | 1.29 ± 0.03 |
| EnHD CM10 | -6.06 ± 0.01 | 4.20 ± 0.01 | -2.30 ± 0.02 | 10 | -0.23 ± 0.01 | -6.23 ± 0.03 | 1.48 ± 0.01 |
| EnHD B3 | -4.15 ± 0.01 | 2.87 ± 0.01 | -0.39 ± 0.01 | 3 | -0.13 ± 0.01 | -4.21 ± 0.02 | 1.46 ± 0.01 |
| EnHD BI10 | -5.08 ± 0.01 | 3.52 ± 0.01 | -1.32 ± 0.01 | 10 | -0.13 ± 0.01 | -4.70 ± 0.04 | 1.34 ± 0.01 |
| EnHD S19 | -7.22 ± 0.01 | 5.00 ± 0.01 | -3.46 ± 0.01 | 19 | -0.18 ± 0.01 | -7.31 ± 0.07 | 1.46 ± 0.01 |
| EnHD SI26 | -7.60 ± 0.02 | 5.27 ± 0.02 | -3.84 ± 0.02 | 26 | -0.15 ± 0.01 | -7.65 ± 0.05 | 1.45 ± 0.01 |
| EnHD SC8 | -5.62 ± 0.10 | 3.89 ± 0.07 | -1.86 ± 0.10 | 8 | -0.23 ± 0.01 | -5.44 ± 0.23 | 1.39 ± 0.05 |
| EnHD WC21 | -6.58 ± 0.06 | 4.56 ± 0.04 | -2.82 ± 0.06 | 21 | -0.13 ± 0.01 | -5.98 ± 0.11 | 1.31 ± 0.02 |
| EnHD K2R | -3.81 ± 0.05 | 2.64 ± 0.03 | -0.05 ± 0.05 | 1 | --- | -3.75 ± 0.03 | 1.42 ± 0.02 |
| EnHD S9T | -4.38 ± 0.04 | 3.04 ± 0.03 | -0.62 ± 0.04 | 1 | --- | -4.14 ± 0.13 | 1.36 ± 0.05 |
| EnHD K17E | -4.57 ± 0.01 | 3.16 ± 0.01 | -0.81 ± 0.01 | 1 | --- | -4.57 ± 0.06 | 1.44 ± 0.02 |
| EnHD L26P | -3.26 ± 0.01 | 2.26 ± 0.01 | 0.50 ± 0.01 | 1 | --- | -3.46 ± 0.02 | 1.52 ± 0.01 |
| EnHD S35A | -4.70 ± 0.01 | 3.25 ± 0.01 | -0.93 ± 0.01 | 1 | --- | -4.32 ± 0.07 | 1.32 ± 0.02 |
| EnHD I45V | -3.24 ± 0.05 | 2.25 ± 0.03 | 0.52 ± 0.05 | 1 | --- | -3.07 ± 0.05 | 1.38 ± 0.01 |
| EnHD I47V | -3.59 ± 0.13 | 2.49 ± 0.09 | 0.17 ± 0.13 | 1 | --- | -3.36 ± 0.08 | 1.36 ± 0.03 |
| EnHD K52R | -4.48 ± 0.09 | 3.10 ± 0.06 | -0.72 ± 0.09 | 1 | --- | -4.65 ± 0.07 | 1.50 ± 0.03 |
| CHD R2K | -8.00 ± 0.02 | 5.53 ± 0.02 | -0.03 ± 0.02 | -1 | --- | -8.64 ± 0.20 | 1.56 ± 0.04 |
| CHD T9S | -7.59 ± 0.01 | 5.26 ± 0.01 | -0.44 ± 0.02 | -1 | --- | -7.94 ± 0.04 | 1.51 ± 0.01 |
| CHD E17K | -7.08 ± 0.01 | 4.90 ± 0.01 | -0.94 ± 0.02 | -1 | --- | -7.20 ± 0.04 | 1.47 ± 0.01 |
| CHD P26L | -7.66 ± 0.01 | 5.31 ± 0.02 | -0.37 ± 0.02 | -1 | --- | -7.60 ± 0.07 | 1.43 ± 0.01 |
| CHD A35S | -6.70 ± 0.04 | 4.64 ± 0.03 | -1.33 ± 0.04 | -1 | --- | -6.53 ± 0.15 | 1.41 ± 0.04 |
| CHD V45I | -7.88 ± 0.03 | 5.46 ± 0.02 | -0.15 ± 0.04 | -1 | --- | -8.67 ± 0.10 | 1.60 ± 0.03 |
| CHD V47I | 8.32 ± 0.01 | 5.76 ± 0.02 | 0.29 ± 0.02 | -1 | --- | -8.87 ± 0.03 | 1.54 ± 0.01 |
| CHD R52K | -7.41 ± 0.01 | 5.13 ± 0.02 | -0.62 ± 0.02 | -1 | --- | -7.39 ± 0.04 | 1.44 ± 0.01 |

[a]Parameters determined from a fits to a global model with a single common *m*-value (1.44 ± 0.01 kcal mol$^{-1}$ M$^{-1}$, where the uncertainty is estimated from the covariance matrix from the fit) but individual $\Delta G^{\circ}_{H2O}$ values for each unfolding transition. Fitted $\Delta G^{\circ}_{H2O}$ values and derived transition midpoints ($C_m$) are means from three independent transitions; uncertainties are standard errors of the mean.

[b]$\Delta\Delta G^{\circ}_{H2O}$ values for single residue variants are determined from globally fitted mean $\Delta G^{\circ}_{H2O}$ values using Eqs 5A and 5B. Uncertainties in $\Delta\Delta G^{\circ}_{H2O}$ values are propagated according to Eq 2.

[c]$\Delta N$ is the number of substitutions toward consensus.

[d]Uncertainties are obtained by dividing uncertainties in $\Delta\Delta G^{\circ}_{H2O}$ values by $\Delta N$.

[e]Parameters determined from local fits of unfolding transitions. Uncertainties represent standard errors of the mean from three unfolding transitions.

**Table S2. Residue side chain relative solvent accessible surface areas in the EnHD crystal structure.**

| Residue | Relative SASA(%)[a] | Classification[b] | Mismatching position[c] |
|---|---|---|---|
| D1 | N/A | S | X |
| K2 | N/A | S | X |
| R3 | 46.7 | I | |
| P4 | 100 | S | X |
| R5 | 38.1 | I | |
| T6 | 42.6 | I | |
| A7 | 100 | S | X |
| F8 | 4.2 | B | |
| S9 | 59.6 | S | X |
| S10 | 100 | S | X |
| E11 | 96.4 | S | |
| Q12 | 21.4 | I | |
| L13 | 39.9 | I | |
| A14 | 80.3 | S | X |
| R15 | 32.6 | I | X |
| L16 | 0 | B | |
| K17 | 62.9 | S | X |
| R18 | 80.8 | S | X |
| E19 | 18.4 | I* | X |
| F20 | 5.9 | B | |
| N21 | 74.5 | S | X |
| E22 | 69.9 | S | X |
| N23 | 52.8 | S | |
| R24 | 46.3 | I | |
| Y25 | 74.6 | S | |
| L26 | 5.4 | B | X |
| T27 | 76.5 | S | X |
| E28 | 91.4 | S | X |
| R29 | 89.1 | S | X |
| R30 | 23.2 | I | X |
| R31 | 19.2 | B | |
| Q32 | 58.4 | S | X |
| Q33 | 65.8 | S | X |
| L34 | 2.5 | B | |
| S35 | 18.4 | B | X |
| S36 | 61.8 | S | X |

| | | | |
|---|---|---|---|
| E37 | 51.6 | S | X |
| L38 | 9 | B | |
| G39 | 82.8 | S | |
| L40 | 5.7 | B | |
| N41 | 53.9 | S | X |
| E42 | 36.7 | I | |
| A43 | 42.2 | I | X |
| Q44 | 6.7 | B | |
| I45 | 0 | B | X |
| K46 | 57.6 | S | |
| I47 | 30.8 | I | X |
| W48 | 1.7 | B | |
| F49 | 1 | B | |
| Q50 | 48 | I | |
| N51 | 47.4 | I | |
| K52 | 18.1 | I* | X |
| R53 | 26.9 | I | |
| A54 | 78.7 | S | |
| K55 | 59.3 | S | X |
| I56 | 47.2 | I | |
| K57 | N/A | S | |
| K58 | N/A | S | |

[a]Residue-specific solvent accessible surface area (SASA) calculations were determined from the EnHD crystal structure (PDB: 1ENH) using GETAREA (main text reference 22). Residues D1, K2, K57, and K58 (denoted with N/A) are not present in the EnHD crystal structure

[b]D1, K2, K57, and K58 were classified as surface residues since they are on the protein N- and C-termini. Residues E19 and K52 show relative SASA values consistent with buried residues, but were classified as intermediate because they are charged residues.

[c]Mismatching residues between EnHD and CHD are noted with an “X.”

**Table S3. Side chain-side chain contacts for sequence differences.**

| Residue | Side chain contacts[a] | Strongly conserved[b] |
|---|---|---|
| D1 | N/A | |
| K2 | N/A | X |
| P4 | | |
| A7 | | |
| S9 | | X |
| S10 | | |
| A14 | | |
| R15 | L34, ***I45***, F49 | |
| K17 | W48 | X |
| R18 | | |
| E19 | ***R30***, L34 | |
| N21 | | |
| E22 | | |
| L26 | L34, ***I45***, F49 | **X** (underlined) |
| T27 | | |
| E28 | | |
| R29 | | |
| R30 | ***E19***, N23, F49 | |
| Q32 | E42 | |
| Q33 | | |
| S35 | E42, ***I45*** | **X** (underlined) |
| S36 | | |
| E37 | ***R15*** | |
| N41 | R3 | |
| A43 | R3 | |
| I45 | ***L26***, L34, ***S35*** | **X** (underlined) |
| I47 | R5 | X |
| K52 | F20 | X |
| I56 | F20, R24 | |

[a]Contacts were counted between two non-hydrogen side-chain atom pairs within 4.2 Å in the EnHD crystal structure (PDB 1ENH) and are separated by more than 5 residues. Residues D1 and K2 are not present in the crystal structure. Residue with no side chain contacts are left blank. Contact partners that are also consensus residues are italicized

[b]The eight strongly conserved sequence differences are noted with an "X." Positions that show non-additivity are underlined.

**Table S4. Residue differences among variants.**

| | EnHD | EnHD SC8 | EnHD WC21 | EnHD CS19 | EnHD CM10 | EnHD B3 | EnHD BI10 | EnHD S19 | EnHD SI26 | CHD |
|---|---|---|---|---|---|---|---|---|---|---|
| **EnHD** | 0 | 8 | 21 | 19 | 10 | 3 | 10 | 19 | 26 | 29 |
| **EnHD SC8** | --- | 0 | 29 | 25 | 4 | 5 | 8 | 21 | 24 | 21 |
| **EnHD WC21** | --- | --- | 0 | 4 | 25 | 24 | 21 | 8 | 5 | 8 |
| **EnHD CS19** | --- | --- | --- | 0 | 29 | 22 | 19 | 10 | 7 | 10 |
| **EnHD CM10** | --- | --- | --- | --- | 0 | 7 | 10 | 19 | 22 | 19 |
| **EnHD B3** | --- | --- | --- | --- | --- | 0 | 7 | 22 | 29 | 26 |
| **EnHD BI10** | --- | --- | --- | --- | --- | --- | 0 | 29 | 22 | 19 |
| **EnHD S19** | --- | --- | --- | --- | --- | --- | --- | 0 | 7 | 10 |
| **EnHD SI26** | --- | --- | --- | --- | --- | --- | --- | --- | 0 | 3 |
| **CHD** | --- | --- | --- | --- | --- | --- | --- | --- | --- | 0 |
